# A common mechanism processes auditory and visual motion

**DOI:** 10.1101/2023.09.05.556424

**Authors:** David Alais, Uxía Fernández Folgueiras, Johahn Leung

## Abstract

We report behavioural findings implying common motion processing for auditory and visual motion. We presented brief translational motion stimuli drifting leftwards or rightwards in the visual or auditory modality at various speeds. Observers made a speed discrimination on each trial, comparing current speed against mean speed (i.e., method of single stimuli). Data were compiled into psychometric functions and means and slopes compared. Slopes between auditory and visual motion were identical, consistent with a common noise source, although mean speed for audition was veridical while visual speeds were significantly underestimated. An inter-trial analysis revealed clear motion priming in both audition and vision (i.e., faster perceived speed after a fast preceding speed, and *vice versa* – a positive serial dependence). Plotting priming as a function of preceding speed revealed the same slope for each modality. We also tested whether motion priming was modality specific. Whether vision preceded audition, or audition preceded vision, a positive serial bias (i.e., priming) was always observed. We conclude a common process underlies auditory and visual motion, and that this explains the closely matched data in vision and audition, as well as the crossmodal data showing equivalent motion priming regardless of the preceding trial’s modality.

## Introduction

Motion perception is usually studied in a single modality although it is clear that the brain must have the capacity to process motion signals from multiple modalities because real objects translating across space often produce correlated motion in more than one sensory modality. A common motion processing network for visual, auditory and tactile motion is a compelling notion because it would be very efficient from an evolutionary view, saving duplication of motion systems in each modality which, in any case, would still need to be integrated to elicit the benefits of multisensory perception^1^. Some authors have argued that the human medial temporal region (hMT+) may be the region that serves this function by responding to motion without regard to the sensory modality of the input^1,2^. hMT+ is a motion-specialised area that is located quite early in the visual cortical pathway, one synapse from primary visual cortex (V1), and is not generally considered to be multisensory. Activity in MT correlates well with perceived visual motion^3-5^ and the movement perceived in the visual motion aftereffect^6-9^.

Human neuroimaging investigations into supramodal motion processing in MT have not produced clear-cut results. The approach has been to test for supramodal motion processing by looking for overlapping responses in hMT+ to auditory and visual motion, or to auditory and tactile motion. Results from these studies are mixed, with fMRI studies testing for auditory motion responses in hMT+ generally producing negative results in sighted subjects^10-12^, and two that did find auditory responses^13,14^ being queried on methodological grounds^15,16^. Other studies have looked for tactile motion responses in hMT+ and found evidence of direction selectivity^17-23^, although again, several of these have been queried methodologically^15^. More recently, Dormal, et al.^24^ used a decoding approach and found that auditory motion direction could be reliably decoded from hMT+ activity. Rezq, et al.^25^ confirmed this decoding result in hMT+ and in an interesting symmetry also showed that response patterns to auditory motion could predict visual motion direction, and visual response patterns could predict auditory motion.

Many behavioural studies have examined audiovisual motion perception but clear evidence of supramodal processing is lacking and there are no reports of optimal integration to parallel those observed for spatial tasks^26,27^. Studies testing for summation of motion signals by comparing bimodal detection thresholds against unimodal thresholds have generally found weak interactions consistent with probability summation rather than an additive or superadditive summation^28,29^. There are many reports showing that sound can modulate visual motion perception^30-32^ but these may be attributable to changes in response criterion rather than sensitivity, and one study^28^ showed small sensitivity benefits regardless of whether the auditory and visual directions corresponded (i.e., an absence of direction selectivity). The study which came closest to finding a linear combination of auditory and visual motion signals^33^ used sound and motion signals that translated horizontally around the observer (we use a similar format in this study) rather than screen-based visual motion drifting within a fixed aperture and a pair of flanking speakers, and they did find a strong summation that only occurred with congruent motion components. A common supramodal process should show bidirectionality between the senses, and to date the only study to report this involved tactile and visual stimuli^34^, finding that adaptation to tactile motion exhibited aftereffects in vision, and *vice versa*.

In the current experiments, we randomly presented various speeds in a trial sequence that was either entirely visual or entirely auditory (Experiment 1) or involved randomly interleaved visual and auditory motions from trial to trial (Experiment 2). To preview the results, Experiment 1 showed speed perception was more veridical in audition than vision, and that there was no difference in the precision of speed discrimination between vision and audition. We also established that both modalities showed motion priming, an attractive effect whereby current speed perception is biased towards the previous trial’s speed (i.e., a positive serial dependence). Experiment 2 randomly interleaved auditory and visual motions moving in leftward or rightward directions. When consecutive trials were congruent in direction (both leftward or both rightward) we found symmetrical cross-modal priming effects between trials. That is, current perceived speed was equivalently primed by the preceding speed, regardless of the modality combination. Pairs of incongruent trials showed no priming. Overall, these results suggest a common and directionally selective mechanism for auditory and visual motion.

## Methods

### Participants and ethics statement

Fifteen participants from the University of Sydney student population took part in the experiment and gave informed consent. All had normal or corrected-to-normal vision and reported normal hearing. The experiment was approved by the Human Research Ethics Committee of the University of Sydney (project number: 2016/662) and procedures conformed to the declaration of Helsinki.

### Apparatus

A PROPixx DLP projector (VPixx Technologies, Canada) was used to project the translating visual stimulus (a vertical Gabor patch) at a framerate of 120 Hz on a white, acoustically transparent projector screen 143 cm away from the subject. The projector had a spatial resolution of 1920 x 1080 pixels and cast a viewable image area 136 x 76 cm in width which corresponded to a visual angle of 57° x 32° and a visual resolution of 35 pixels per degree. The projector was connected via a DataPixx2 display driver and controlled using software written in MATLAB 2014a (The MathWorks, Inc.) running on an Apple Mac Pro computer and had a linearised luminance output. Participants gave their responses using a ResponsePixx (VPixx Technologies Inc.) button box.

A custom-made soundbar, consisting of 11 speakers (Visaton F8SC) fixed horizontally on a wooden support and aligned with the visual motion path, was used to deliver the moving auditory stimulus in free field. The soundbar was mounted on a wall 150 cm away from the subject at eye level and the linear separation of each speaker was fixed at 5° relative to the subject’s head (see Fig. 1). These speakers were controlled by two RME Fireface 400 audio interfaces (RME Audio) and were hidden from view by the large (169 x 100 cm) acoustically transparent projector screen sitting 7 cm in front of the soundbar.

**Figure 1.**
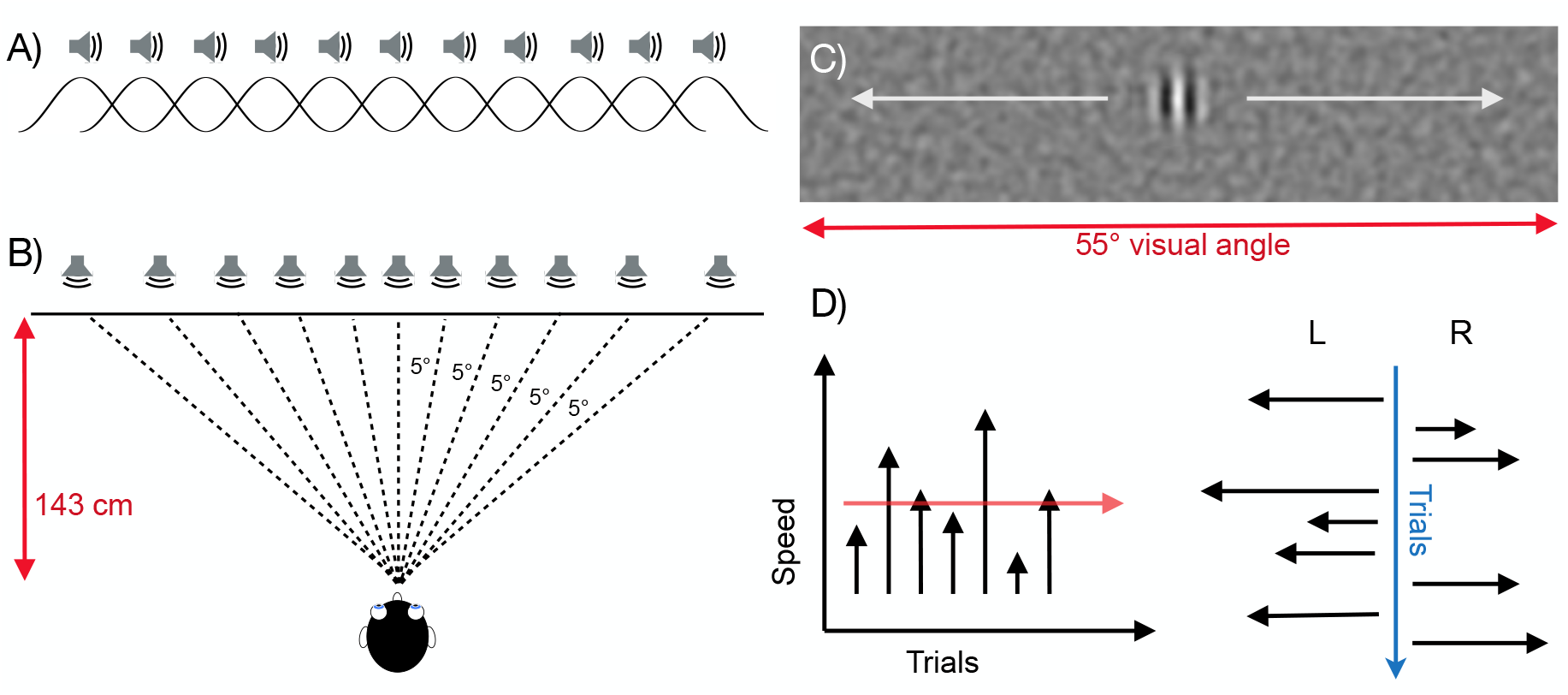
Stimuli and methods. (A) Auditory motion was produced by crossfading a sound signal between a series of horizontally aligned speaker locations to produce smooth continuous movement over space. Crossfading in this way varies inter-aural time and level differences as well as spectral cues to produce a strong motion signal. The signal was an amplitude-modulated white noise, with a modulation frequency of 20 Hz and modulation depth of 0.2. (B) Speaker locations were spaced to have a constant spatial separation of 5° of lateral angle from the participant’s position. (C) The visual stimulus was a high-contrast vertically oriented grating (0.8 cyc/deg) in a Gaussian window which translated horizontally over a background of low-contrast dynamic noise that was spatially filtered into a passband centred on the grating’s frequency. The stimulus was projected onto an acoustically transparent screen placed in front of the speakers so that they were not visible. (D) Seven speeds were randomly interleaved (20, 40, 50, 60, 70, 80, 100 deg/sec) and participants judged whether a given speed was faster or slower than the mean of the set^37,38^, as represented in the left panel where the red arrow illustrates the mean. Over trials, leftward and rightward motions were interleaved randomly (right panel). To encourage reliance on velocity, path length and duration were randomly jittered over trials.

### Stimuli

The visual motion stimulus was a vertically oriented Gabor patch (Gaussian standard deviation = 0.8°; grating spatial frequency = 0.8 cyc/deg at 0.75 of maximum contrast) that translated horizontally across the screen on a background of spatially filtered noise with centre frequency of 1.0 cyc/deg, a one-octave passband and a contrast of 0.25 of maximum. To make total duration and the start/end points unreliable as cues for judging speed, the motion path length varied from trial to trial by randomly jittering the start and end points. For the auditory stimulus, this was done by randomly selecting the start point on each trial from speaker 1, 2 or 3 and the end point from speaker 9, 10 or 11. For the visual stimulus, path lengths varied randomly in a continuous range between 30 and 40 degrees. The location of the translating Gabor was randomly jittered within a 20-pixel range on every frame but maintained a constant average speed. For both stimuli, once path length was selected, trial duration was then varied to obtain the required speed.

### Auditory Motion

The auditory stimulus consisted of an amplitude modulated (*F*c = 20 Hz) broadband white noise sampled at 48 kHz that was created at the beginning of each trial. This noise was divided into 11 segments, one for each of the 11 speakers along the soundbar. Smooth apparent motion was created by playing each speaker in sequence using a crossfade and by applying a sine window to the onset and offset of each noise segment. To eliminate loudness variations between individual speakers, the sound pressure level of each speaker was tested and calibrated to ensure equal loudness of (∼75 dBA), as measured at a distance of 1 m with a sound pressure meter. To prevent subjects from using distance and location cues as a basis for velocity judgements, the start- and end-points of the sounds were randomly jittered by ±15° (3 speaker intervals, see Freeman et al, 2014). All stimuli were generated on a MacPro (Apple Inc.) running MATLAB (R2016b, The MathWorks, Inc., Natick, Massachusetts, United States) and Psychophysics Toolbox extensions^35,36^.

### Design and procedure

Participants did a speed discrimination task in a self-paced experiment using the “method of single stimulus”^39^. In the unimodal conditions, a speed drawn from a set of seven (20, 40, 50, 60, 70, 80 & 100°/s) was presented and subjects were asked to report whether they perceived the speed as faster or slower than the average of all previously presented speeds. This paradigm has been used in previous speed discrimination studies^37^ to determine the point of subjective equality (PSE) and discrimination threshold and requires only half as much stimulus exposure as two-alternative forced-choice methods and produces better discrimination thresholds^38^. In effect, the method is a two-alternative comparison involving the current stimulus and an internal standard of mean speed.

We examined unimodal and crossmodal conditions separately in two sessions over two days in a counterbalanced order. In the unimodal conditions, every trial in an experimental block consisted of only auditory or visual motion. The seven velocities from the stimulus set were tested for both leftward and rightward directions, each repeated 20 times for a total of 280 trials per block. Subjects completed two blocks for each modality, alternating between audition and vision, and took breaks between each block. The order of presentation within a block (direction and speed) was completely randomised, as shown in Figure 1d, so that direction was unpredictable. This meant that consecutive directions could occur in the same direction (congruent) or opposite direction (incongruent). When analysing unimodal motion priming (Figure 3), only congruent data was used.

**Figure 2.**
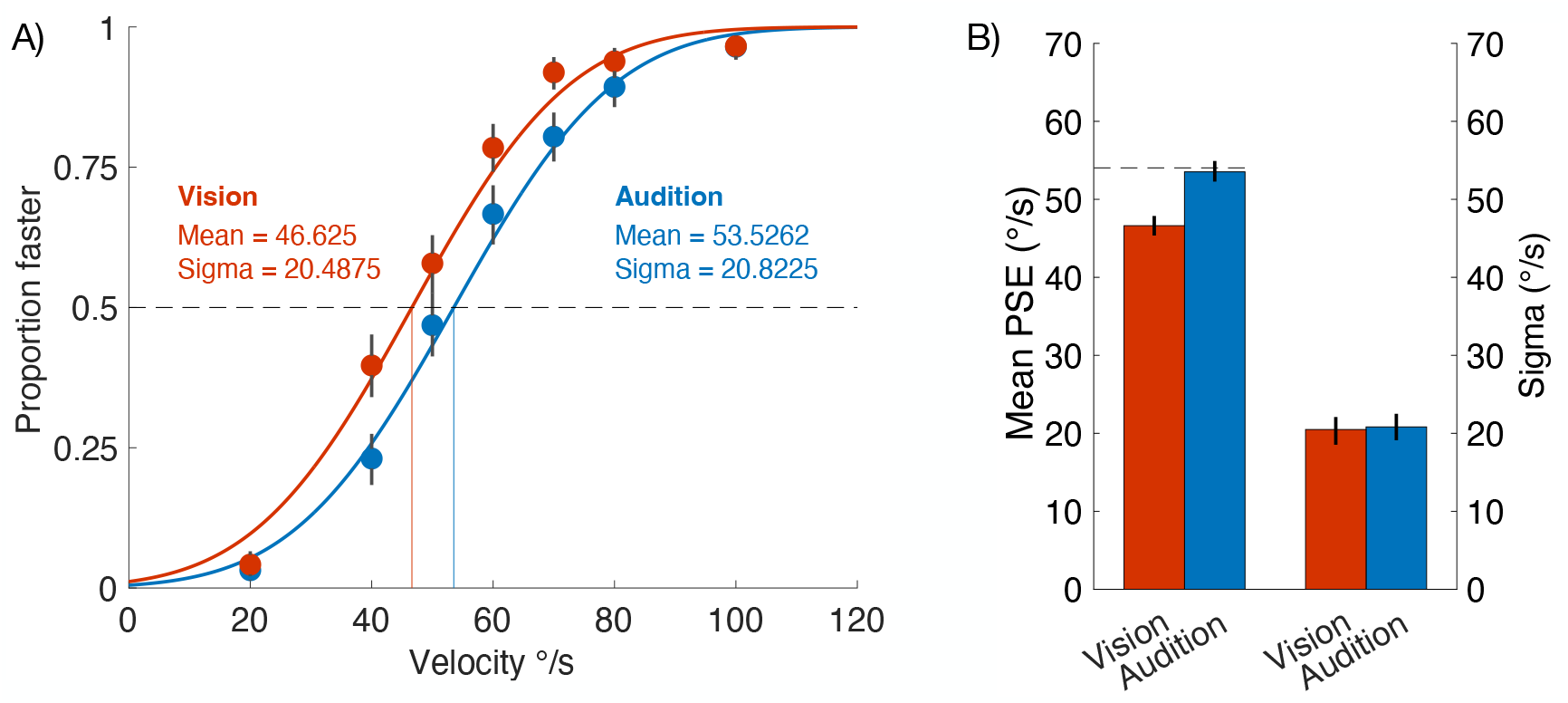
Unimodal data showing velocity discrimination of a set of velocities around a standard velocity with a geometric mean of 54.03 °/s. (A) The data points show mean ‘proportion faster’ for each level of velocity for all observers pooled into a single super subject and error bars show 95% confidence intervals produced by bootstrapping the data 10,000 times. The continuous lines show the best-fitting cumulative Gaussian psychometric function. (B) Parameters from the best-fitting psychometric functions for audition and vision. The means (left) and bandwidth (right) are plotted with errors bars showing 95% confidence intervals from 10,000 bootstraps. The dashed line shows the geometric mean speed.

**Figure 3.**
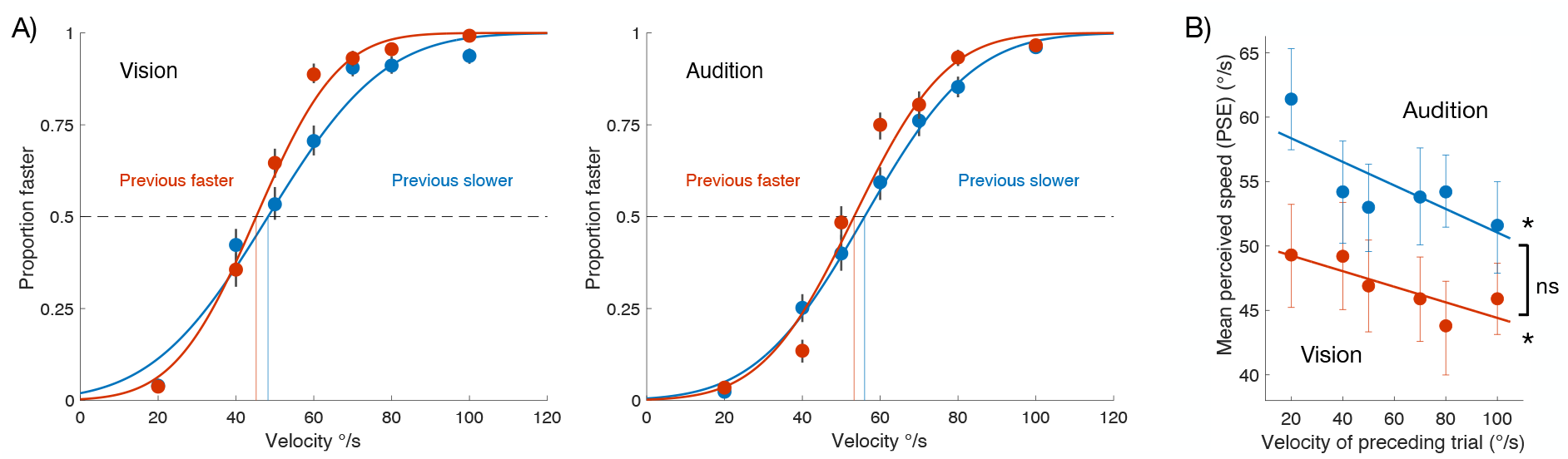
Unimodal data showing significant motion priming of the current trial’s perceived velocity by the previous trial’s velocity. (A) For each modality, trials were binned into two groups based on the speed of the preceding trial (‘previous slower’ = 20, 40, 50 °/s; ‘previous faster’ = 70, 80, 100 °/s) and psychometric functions were fitted to each group. For both modalities, the PSE for the ‘previous faster’ group was significantly lower than for the ‘previous slower’ group. Error bars show ±1 standard deviation based on 10,000 bootstraps. (B) Motion priming for all levels of previous velocity. PSEs decline (more “perceived faster” responses) as preceding velocity increases. Bootstrap sign-tests confirmed the slopes of the best-fitting lines for audition and vision were both significantly negative but did not differ from each other (p = .2078). Error bars show 95% confidence intervals based on 10,000 bootstraps.

In the crossmodal condition, auditory and visual trials were interleaved in a balanced alternation (either A, V, A, V … or V, A, V, A …). The crossmodal condition contained no bimodal trials, only interleaved unimodal trials, and subjects again reported whether the current stimulus moved faster or slower than the average of the set of all stimuli. The velocities tested were as above (20, 40, 50, 70, 80 & 100°/s), in leftward and rightward directions, each repeated 10 times per block (240 trials) and subjects completed two blocks with a break in between. Again, direction of motion (leftward or rightward) was randomised to ensure unpredictability. When analysing crossmodal motion priming (A, V or V, A) we analysed separately for congruent priming (same direction: Figure 4) or incongruent (opposite directions: Figure 5).

**Figure 4.**
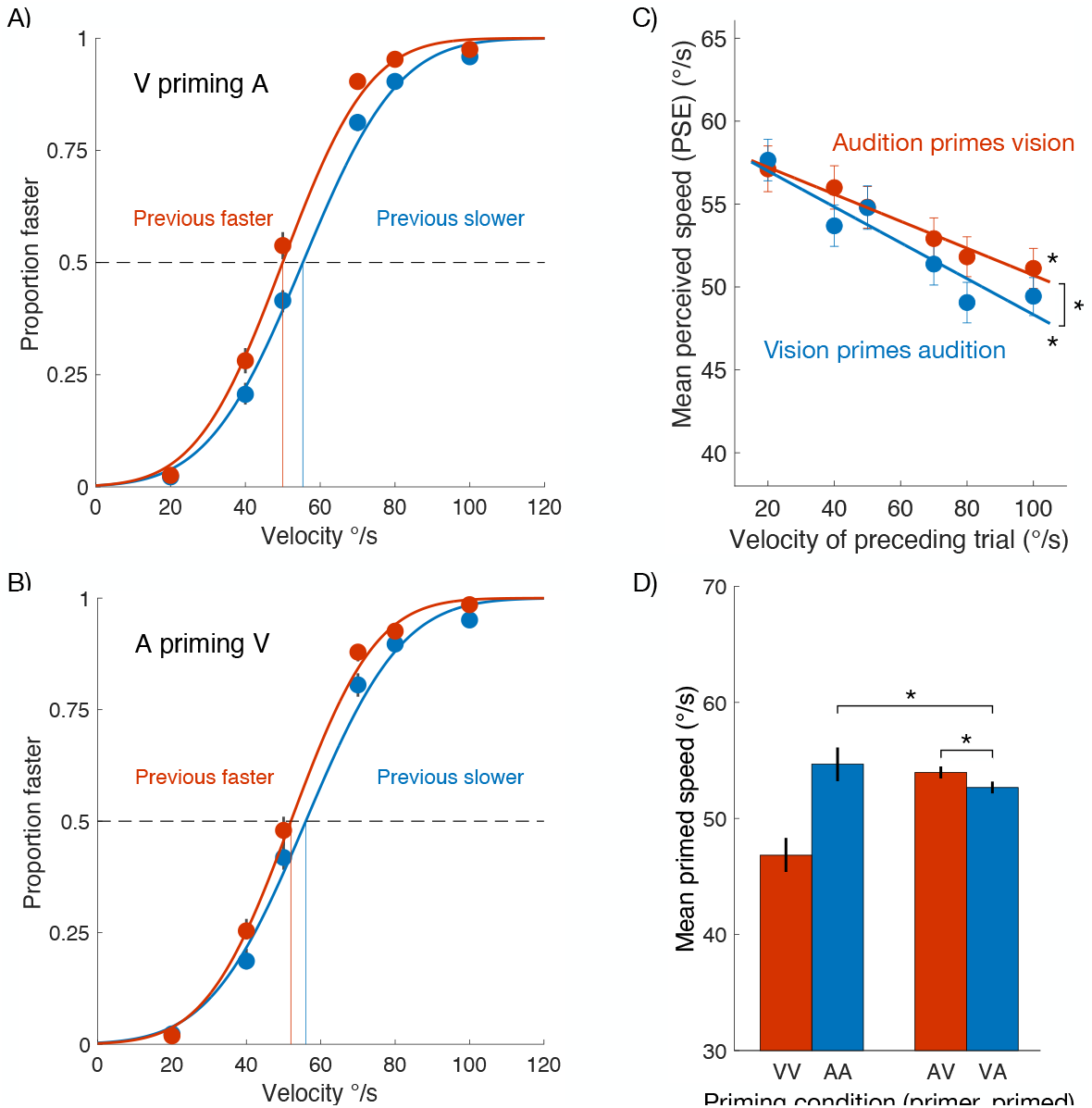
Motion priming between modalities with congruent directions. (A) Auditory motion primes visual motion (AV). Data for judging visual speed were binned into ‘previous slower’ [20, 40, 50 °/s] and ‘previous faster’ [70, 80, 100 °/s] groups and psychometric functions fitted to each group. There was a significant AV priming effect: the PSE was significantly lower after ‘previous fast’ trials (p < .0001). (B) Visual motion primes auditory motion (VA). Same format as panel A, except auditory motion preceded visual motion and produced a significant priming effect (p < .0001). (C) Motion priming for all levels of previous velocity. PSEs decline (more “perceived faster” responses) as preceding velocity increases for both AV and VA conditions. A bootstrap sign-test indicated the slopes of the best fitting lines differed significantly between the conditions, with VA having a steeper slope than AV (p = .0009). (D) Columns on the right show the mean crossmodal priming effect from panel C. For comparison, the means of the unimodal priming functions from Figure 3b are shown. Error bars in all panels are 95% confidence intervals based on 10,000 bootstraps.

## Results

### Speed discrimination for vision and audition

Data from the visual and auditory unimodal conditions were analysed separately and are shown in Figure 2. The data for leftward and rightward directions were very similar and did not differ significantly and were therefore pooled. For each subject, trials were sorted by stimulus speed and mean ‘proportion faster’ judgements were calculated for each level of speed. The mean proportions at each speed for all observers were pooled into a super subject analysis and a cumulative Gaussian psychometric function was fitted. The data were bootstrapped 10,000 times to compute 95% confidence intervals. The mean of the best-fitting cumulative Gaussian was taken as the point of subjective equality (PSE) and the Gaussian’s standard deviation (sigma) provided the speed discrimination threshold. Figure 2a shows the visual and auditory data overlaid for comparison. Data for both modalities show the expected pattern of ‘proportion faster’ judgements rising as stimulus speed increases, confirming that the method of single stimuli used here yields reliable and orderly data for auditory motion discrimination – as previously shown for visual motion^37,38^.

Figure 2b compares the PSEs and discrimination thresholds for auditory and visual motion. The PSE effectively indicates the mean perceived speed of the stimulus set and for comparison the dashed horizontal line in 2b shows the geometric mean of the set of speeds used (54.03°/s). While the arithmetic mean speed is 60°/s, velocity is represented logarithmically in cortex (Nover, Anderson & DeAngelis, 2005) and thus the geometric mean is more appropriate. As the confidence intervals show, the PSE for vision (46.625°/s; CI_95_ = [45.196–47.909]) was significantly lower than the mean stimulus speed and was also significantly lower than the PSE for audition (53.526°/s; CI_95_ = [52.133–54.867]). This represents a reliable underestimation of speed in vision relative to audition. Moreover, mean perceived speed for auditory motion was surprisingly veridical and did not differ from the geometric mean speed. An interesting feature of the unimodal psychometric functions is the closely matched discrimination thresholds for auditory and visual motion. Motion discrimination is generally considered poor for audition^40^ yet here the speed discrimination thresholds (i.e., the bandwidth or standard deviation of the psychometric function) for vision (20.488°/s; CI_95_ = [18.501– 22.295]) and audition (20.823°/s; CI_95_ = [19.174–22.666]) were very similar and not significantly different.

### Inter-trial priming of speed discrimination within modalities: vision vs audition

We investigated whether there were any inter-trial dependencies in our stimulus sequences. Even though our stimulus presentations were brief, a given trial in a sequence is often primed by the preceding one such that current percept shows a bias towards the previous one. This is a priming or ‘positive serial dependence’ effect that occurs for a wide range of stimuli^41-44^ including motion^45^. We conducted a sequential dependency analysis which involved binning all trials into two groups based on whether the speed on the preceding trial was slow (20, 40 & 50°/s) or fast (70, 80 & 100°/s). For each group, we then calculated the best-fitting psychometric functions and compared the PSEs. If the previous trial has no influence on the current one, then all trials would be independent and the psychometric functions calculated from the ‘previous slow’ and ‘previous fast’ groups should not differ in PSE.

Figure 3 shows the results of our serial dependence analysis. For both audition and vision, rates of ‘proportion faster’ responses were higher when the previous trial was fast, compared to when the previous trial was slow (Fig. 3A). This pattern is consistent with motion priming, a positive (or ‘attractive’) serial dependence effect often seen when preceding motion stimuli are brief^46-48^ and contrasts with motion adaptation – a negative (or ‘repulsive’) serial dependence that requires longer adaptation periods and produces the classic motion aftereffect^49,50^. A bootstrap sign-test confirmed that PSEs were significantly lower (leftward shift of the psychometric function) after ‘previous fast’ trials for both vision and audition (vision: p < .0001; audition: p < .0001), establishing that the previous trial was sufficient to prime the current trial and that vision and audition show the same effect.

With the priming effect established at the level of fast vs slow, we went further and analysed the PSE shift more finely. To do this, we split the ‘previous slower’ group into three smaller groups corresponding to its three levels of previous velocity (i.e., 20, 40, 50 °/s) and fitted psychometric functions to each of the groups. We did the same for the ‘previous larger’ data (splitting it into three previous velocities: 70, 80, 100°/s). Figure 3b shows the PSEs as a function of these previous velocities, for both audition and vision. Dividing the data in this way reduces the observations in each group by a factor of three yet there is still a clear and significant negative slope for each modality (audition: slope = -.0918, p = .0008; vision: slope = -.0594, p = .0181), thus showing the same priming relationship as in Figure 3A (previous faster speeds consistently shifting PSEs to the left, indicating increasing levels of “perceived faster” responses). The priming functions for auditory and visual motion are very similar. To test if they were different, we bootstrapped the data for each level of previous speed 10,000 times and re-calculated the best linear fit across the previous speed levels 10,000 times. If the auditory and visual data have different slopes, then the differences between the bootstrapped slopes for audition and vision should be consistently different from zero. A bootstrap sign-test confirmed this was not the case (p = .2078). Thus, motion priming effects as a function velocity are very similar for both auditory and visual motion.

### Crossmodal motion priming

We next tested for an effect of motion priming between vision and audition by examining the data from the crossmodal condition in which auditory and visual motions were presented in alternation (A,V or V,A) over trials with direction randomly determined on each trial (left or right). For the analysis shown in Figure 4, we examined only pairs of trials that had congruent directions. This design allowed us to test for crossmodal motion priming in both directions – audition priming vision (AV), and vision priming audition (VA)and the results are shown Figure 4. The psychometric functions in Figure 4a show auditory speed judgements when preceded by either a fast (red) or a slow (blue) visual motion. Figure 4b shows the converse: visual speed judgements when preceded by fast or slow auditory motion. As observed within modalities, we found significant motion priming between modalities in both directions. A bootstrap sign-test based on 10,000 iterations confirmed that PSEs were significantly lower (leftward shift of the psychometric function) after ‘previous fast’ trials for both vision priming audition (VA: p < .0001) and for audition priming vision (AV: p < .0001). This demonstrates that the previous trial does not need to be in the same modality to prime a subsequent motion, and that the motion priming effect occurs symmetrically, with the AV and VA conditions producing equivalent results.

We divided the ‘fast’ and ‘slow’ data further to examine the priming effect as a function of each level of prime speed. Figure 4c plots the PSEs from the psychometric functions computed for each level of preceding motion. For both the AV and VA conditions, motion priming shows a clear linear dependence on the speed of the priming stimulus that is well described by a linear function. The results mimic those in the Figure 3b and clearly illustrate that motion priming also occurs for alternating auditory and visual trials. Again, we bootstrapped the data for each preceding velocity 10,000 times and fitted linear functions across prime speed for each iteration to compare slopes for the AV and VA condition. The slopes were significant for each modality order (AV: slope = -.0763, p < .0001; VA: slope = -.1249, p < .0001) and quite similar, although the difference was statistically reliable (p = .0009) with VA priming showing a stronger effect than AV.

Finally, we averaged the PSEs in the AV and VA priming functions in panel C into a single value (Figure 4d, right) and compared them with the average of the unimodal priming functions plotted in Figure 3c (see Figure 4d, left). This shows whether the PSE (i.e., mean perceived speed) for a given modality differs depending on whether it was primed by the same or different modality. The biggest difference occurs for the visual PSE (Figure 4d, red columns): when primed by visual motion (VV), the mean PSE was 46.843°/s and rose to 52.715 (+5.872°/s, p < .0001) when primed by auditory motion (AV). A smaller and opposite effect occurred for the auditory PSEs (blue columns): when primed by auditory motion (AA), the mean PSE was 54.717°/s and fell slightly to 54.091 (-0.626°/s, p = .8656) when primed by visual motion (VA).

## Discussion

In these experiments we measured speed discrimination for a range of horizontally translating auditory and visual motions. Using the same set of speeds in each modality (20, 40, 50, 60, 70, 80, 100°/s) and employing the method of single stimuli in which the speed on a given trial is compared against the mean of the set of speeds (see^38,39^), we first compared vision and audition in unimodal experiments. The unimodal data revealed that the slopes of the psychometric functions for auditory and visual motion were the same, although their means differed significantly. The mean of the auditory motion psychometric function matched the mean of the stimulus set, while the mean of the visual function was significantly lower than the mean stimulus speed. We also examined motion priming, that is, the tendency for a current motion speed to be biased towards the speed of the preceding one, in both unimodal and crossmodal experiments. These analyses showed a reliable unimodal motion priming effect occurred and that an equivalent effect occurred crossmodally when the prime stimulus was in the other modality. Taken together, the matching psychometric slopes for auditory and visual motion, and the presence of motion priming, regardless of whether the prime stimulus was in the same or different modality, suggest a common process underlies auditory and visual speed discrimination.

A key finding supporting common processing of auditory and visual speed is the near-identical bandwidths of the cumulative Gaussian psychometric functions for auditory and visual motion shown in Figure 2b. The width of a psychometric function is indicative of the noise associated with the underlying perceptual process and thus with perceptual precision. A narrower bandwidth (i.e., steeper slope) indicates greater precision. The finding of equal precision for auditory and visual speed discrimination is somewhat surprising because motion perception in audition is typically found to be much poorer than in vision^40,51,52^. Weber fractions for visual speed discrimination in vision, for speeds similar to the lower part of the range used here, are on the order of 0.05^53,54^ whereas Weber fractions for equivalent auditory speeds (30°/s & 60°/s) are many times higher at 0.24 and 0.30, respectively^55^. The reason for this large difference is not clear but one key point differentiating the stimuli used in these studies may be relevant. In audition, the stimulus was a sound source that translated across space from one point to the another, like an auditory object. There is no such ‘object displacement’ in vision, as the motion stimuli were a continuous drift within an aperture. It is thus locally contained and does not actually go anywhere. For this reason, comparing these auditory and visual motion discrimination studies is not a like for like comparison. Our study matched vision and audition by using translation across space and the same paths in both modalities (see Fig. 1). Our conditions, therefore, are directly comparable and clearly reveal a strikingly similar psychometric slope for motion discrimination in each modalities.

The difference in the psychometric means between audition and vision was another interesting result. As noted in the previous paragraph, auditory motion is often regarded as inferior to visual motion based on data showing poorer speed discrimination and on the grounds that audition – unlike vision – is fundamentally not a spatial sense. And yet, our data show that it was auditory motion that was veridically perceived. The auditory mean in Figure 2b (blue column) was not significantly different from mean stimulus speed, while the visual mean by contrast was significantly below the stimulus mean, showing an overall underestimation of perceived speed. Again, the reason for this is not clear as visual speed experiments are not done with translating objects as we have used here and we have not been able to find any other studies using a ‘spatial displacement’ approach comparable to ours. One relevant reason for the result, however, might be the nature of the speed cues in each modality. In vision, motion processing begins in primary visual cortex where direction-selective cells with small receptive fields are found. These are pooled into MT cells with much larger receptive fields and a higher range of speed tunings^4,56,57^. It is possible that by using a relatively fast set of speeds (up to 100°/s) there was a degree of smearing at the local V1 level such that MT inherited impaired motion signals. By contrast, the auditory system uses different cues to detect horizontal motion based primarily on interaural time and level differences, supplemented with spectral cues and doppler cues^40,58^ and these are not tied to small spatial receptive fields in the way that vision is and may remain veridical at higher speeds.

The final point of interest comes from the motion priming results. As is well established in the visual domain^46-48^, a brief preceding motion causes the current motion stimulus to be biased towards the preceding one in speed or direction (i.e., motion priming) while a long preceding motion causes a bias away from the preceding one (i.e., motion aftereffect). As our stimuli were brief, we expected and obtained significant motion priming effects for visual stimuli (Fig.3a). We could find no studies of motion priming with auditory stimuli on which to base a prediction, but in the event we also obtained a very similar motion priming effect for auditory motion (Fig. 3b,c). This outcome was far from certain as auditory motion involves a completely different set of cues to visual motion but was indicative that there may be common motion processing for visual and auditory stimuli. The critical condition was the crossmodal case, where auditory and visual stimuli were interleaved. As shown in Figure 4a,b, we obtained clear crossmodal motion priming. That is, consistent with a common motion process, current auditory motion speed was primed by the preceding visual speed, and *vice versa*, and both effects showed very similar functions when plotted for each level of preceding speed (Fig. 4c).

A further interesting feature is shown in Figure 4d, which shows the means of the crossmodal psychometric functions together with the unimodal means from Figure 2. Recall that the mean speeds for unimodal vison and audition were significantly different, with visual stimuli being perceived as slower than auditory (Fig. 2b). On priming trials where vision was preceded by audition, the psychometric mean speed for vision increased relative to the unimodal mean. Conversely, on auditory trials preceded by vision, the psychometric mean speed for audition decreased relative to unimodal audition. In both cases, there was a kind of averaging across the modalities of the current and previous speeds. If each modality was processed independently, the AV trials should have the same mean speed as the unimodal V trials, as AV are actually vision trials (preceded by audition). Similarly, independence would predict VA trials to have the same mean speed as unimodal A trials. Instead, in both crossmodal conditions, mean speed reflected a mix of both component speeds, consistent with common processing.

Visual motion processing is very well understood, first appearing in primary visual cortex (V1) and very strongly present in subsequent areas MT and MST (hMT+). Robust evidence that area MT/hMT+ is critical to speed perception comes from several neurophysiological studies in nonhuman primates. In macacque area MT, neurons are selectively tuned to speed^56,57^ and lesions to MT impair speed discrimination^4,5,59^. In addition, when speed is misperceived, it can still be accounted for by MT responses^3^. Complementing this, work in human neuroimaging shows that speed discrimination preferentially activates hMT+^60^. By contrast, evidence for motion selectivity in primary auditory cortex is scant^11,61^ although several studies have found evidence for auditory motion selectivity in the planum temporale, an area downstream from A1 located posterior to it. PT is primarily involved with language and music, both of which involve motion over the frequency dimension, but several studies have shown it also exhibits robust responses to auditory movement over space^25,62-66^.

The nature of auditory motion processing is still debated. Selectivity for movement over space is not a key feature of the auditory system as the primary auditory representation is tonotopic rather than spatial. For a long time, the standard model has been the snapshot model, with auditory motion inferred from a series of two or more static samples or ‘snapshots’^40,52,67^. Consistent with this position, psychophysical data shows that distance and duration are the strongest cues determining auditory motion perception^68^. A recent model has added further nuance to the snapshot theory by adding a simple temporal integration period and finding it adds further explanatory power^58^. The continued viability of the snapshot model raises the possibility that evidence appearing to indicate motion selectivity in neurophysiological and neuroimaging studies^64,66,69^ could instead arise from samples of positional information along a motion trajectory. The notion of computing motion from a series of well-spaced static positions is very well known in vision where it is known as long-range apparent motion^70^ and such motion stimuli are very effective at activating the visual system’s specialised motion area, hMT+^71-73^.

A number of studies have tested for a shared cortical representation of auditory and visual motion, in both PT and hMT+. Alink, et al.^62^ found reliable PT activation for auditory motion and also found an occipital area from which auditory direction could be decoded – although it was located ventrolaterally from the typical hMT+ location. More recently, Rezk, et al.^25^ found auditory and visual motion were both represented in right hMT+ and further showed that, in right hMT+, responses from motion in one modality could successfully decode motion from the other. This study establishes a direction-selective representation for auditory and visual motion in the same area. Previous studies have also reported that tactile motion also actives hMT+^19,23^, suggesting that it might be effectively a supramodal motion area. In the case of auditory motion, there is evidence from functional and diffusional MRI in human subjects indicating white matter connections between PT and hMT+^74^. This connectivity is a plausible pathway allowing auditory motion signals to activate hMT+.

Overall, we find clear evidence of common processing of auditory and visual motion in a speed discrimination task. Our findings fit well with an emerging picture from neuroimaging studies showing that auditory and visual motion are both represented in hMT+^25^. While visual motion is encoded earlier in the visual pathway in V1 neurons, it is activity in hMT+ that correlates with the perception of motion, as discussed above. If motion from both modalities were processed in hMT+, it would parsimoniously explain the unimodal observations of near-identical psychometric slopes and very similar priming functions, as well as the crossmodal data showing priming between vision and audition in both directions. Because hMT+ responds very effectively long-range apparent motion (a series of position ‘snapshots’^71^), proposing hMT+ as the region of common audiovisual motion processing is agnostic about the ultimate nature of auditory motion as it can be driven by smooth continuous motion or by snapshots of discrete positions over time.

